# Incorporating continuous characters in joint estimation of dicynodont phylogeny

**DOI:** 10.1101/2025.03.03.641281

**Authors:** Brenen M Wynd, Basanta Khakurel, Christian F Kammerer, Peter J Wagner, April M Wright

**Affiliations:** Department of Biological Sciences, Southeastern Louisiana University, Hammond, Louisiana, USA; GeoBio-Center, Ludwig-Maximilians-Universität München, 80333 Munich, Germany; Department of Earth and Environmental Sciences, Paleontology & Geobiology, Ludwig-Maximilians-Universität München, 80333 Munich, Germany; North Carolina Museum of Natural Sciences, Raleigh, North Carolina, USA; Department of Biological Sciences, North Carolina State University, Raleigh, North Carolina, USA; Dept. of Earth & Atmospheric Sciences, and School of Biological Sciences, University of Nebraska, Lincoln, Nebraska, USA

**Keywords:** Continuous traits, Discrete traits, Phylogenetic methods, Morphological data, Bayesian phylogenetics, RevBayes

## Abstract

Continuous characters have received comparatively little attention in Bayesian phylogenetic estimation. This is predominantly because they cannot be modeled by a standard phylogenetic Q-matrix approach due to their non-discrete nature. In this paper, we explore the use of continuous traits under two Brownian motion models to estimate a phylogenetic tree for Dicynodontia, a well-studied group of early synapsids (stem mammals) in which both discrete and continuous characters have been extensively used in parsimony-based tree reconstruction. We examine the differences in phylogenetic signal between a continuous trait partition, a discrete trait partition, and a joint analysis with both types of characters. We find that continuous and discrete traits contribute substantially different signal to the analysis, even when other parts of the model (clock and tree) are held constant. Tree topologies resulting from the new analyses differ strongly from the established phylogeny for dicynodonts, highlighting continued difficulty in incorporating truly continuous data in a Bayesian phylogenetic framework.

## 1 Introduction

Phylogenetics strives to use organismal data that reflect common inheritance and are directly comparable across taxa in the sample set. In working with extant taxa, molecular information (DNA, RNA, amino acid sequences) can offer objective measures that can be compared across species. A single nucleotide base is generally assumed to have the same chemical structure and properties as all other nucleotides of the same type in an alignment or across species. The same is not true for anatomical data (Lewis, 2001; Scotland et al., 2003; Huelsenbeck and Rannala, 2003; Brazeau, 2011; Klopfstein et al., 2015; Wright et al., 2016; Simões et al., 2023; Khakurel et al., 2024), for which structure and expression is often unique not only for individual taxa, but is also filtered through human perception and interpretation and thus varies between individual researchers. Morphological character states almost never reflect truly identical structures everywhere they appear in a character matrix (Lewis, 2001), and different authors may code similar structures differently (Jardine, 1969). Even among presence/absence characters, though “absent” is (on one level) objectively comparable, “present” often represents different and not necessarily equivalent additions to the organisms’ morphology. These types of characters are ubiquitous: across sampled morphological datasets, roughly 70% of characters are binary in this manner (Barido-Sottani et al., 2020). Multistate discrete characters represent their own difficulties in terms of repeatability. State spaces may be misspecified, leading to incorrect homology statements and model parameterization (Hoyal Cuthill, 2015; Khakurel et al., 2024; Mulvey et al., 2024).

Historically, discrete morphological characters are heavily represented in fields that lack ready access to DNA (e.g., paleontology) because of their utility in parsimony-based cladistics (Wright and Wynd, 2024), a tree-estimation criterion that seeks to generate the most likely tree by finding the fewest number of transitions possible among characters (i.e., Occam’s Razor). Parsimony has been the dominant phylogenetic tool for paleontologists. With the growing usage of Bayesian methods in morphological phylogeny estimation, Bayesian estimation methods have been evaluated for discrete characters (Lewis, 2001; Wright and Hillis, 2014; O’Reilly et al., 2016; Klopfstein et al., 2015; Wright et al., 2016; Klopfstein et al., 2019). Because of their ready availability from prior discrete datasets intended for parsimony-based analyses, authors have been able to evaluate these methods using both simulated and empirical datasets. Though the methods themselves have been evaluated, there are still considerable questions surrounding how morphological data should be coded for Bayesian phylogenetic analysis. The fundamental units (such as ‘0’ or ‘1’) are not necessarily comparable across a data matrix (Pogue and Mickevich, 1990; Lewis, 2001; Nylander et al., 2004). For example, a muscle scar having a relatively larger attachment area may be considered, *a priori*, to be faster evolving or ‘easier’ to evolve than the introduction of a brand new muscle scar, but in a parsimony framework, they would both be scored as ‘present’ (1) and would be analytically indistinguishable from one another. How discretizing data in matrix assembly, and what information is lost in doing so, affects overall estimation is still an open question.

Discrete morphological characters often seek to represent complex and difficult-to-measure traits as binary or multistate characters, with numeric states corresponding to a more precise character definition. The alternative to discretization is quantifying traits via direct measurement of dimensions or proportions into continuous variables. Ratios are another type of continuous trait, and tend to be favored in some circumstances as raw measurements are more likely to be influenced by ontogeny when a sample includes differently sized individuals (Bardin et al., 2017; Napoli, 2024; Griffin et al., 2021). Direct measurements of anatomical features, such as lengths or angulations of structures or ratios of one structure to another, have the potential to be more objective than qualitative character state assignments, although worker choice in what features to measure remains a source of some bias. Continuous trait use in phylogenetics has been controversial over the years, with some authors arguing that continuous traits are too noisy for phylogenetics (Chappill, 1989), while others argue that they are the “Holy Grail” of phylogenetic characters (Thiele, 1993). In recent years, there has been a renewed interest in continuous characters from biology and anthropology (Parins-Fukuchi, 2018; Zhang et al., 2024; Matzig et al., 2024). Continuous variables are difficult to model in phylogenetics, as the degree of change over a single unit of time can be infinitesimal and thus are ill-equipped for the transition matrices that form the basis of both cladistic and likelihood-based frameworks. Perhaps as a result of both issues, continuous characters are comparatively rare in the phylogenetic literature. For example, in Graeme Lloyd’s repository of character matrices (Wright et al., 2016), a commonly-used database of morphological datasets, only 10 out of 573 matrices have any continuous traits. The others rely solely on discrete characters.

Many of the objections to and difficulties in using continuous anatomical measurements in phylogenetic analyses are predicated upon using minimum-steps parsimony methods to infer phylogeny, and thus might not pertain to likelihood and Bayesian analyses. While a continuous traits constitute a minority of datasets, parsimony-based cladistics is able to incorporate continuous variables via the additive method (Farris, 1970), which samples sister taxon relationships from a given character and seeks to minimize the difference between nodes moreso than tips. Conceptually, the additive method can be thought of as an inversion of Felsenstein’s independent contrasts (Felsenstein, 1985a). In an additive model, trait space cannot be explored by jumping over intermediate states (Goloboff et al., 2006). The additive model for continuous characters, while not explicitly modeled as such, essentially follows a single rate Brownian motion model (i.e., random walk) for evolution (Felsenstein, 1985b), where the intermediate states are dependent on the terminal states, such that the model retains memory (Goloboff et al., 2006). However, instead of the amount of change per unit time being fixed to a specific rate (*σ*) that applies to branch lengths as in a Brownian motion model, it was instead drawn from a distribution formed by the range of tip states for the descendants of the node in question (Farris, 1970). Although the philosophies behind Brownian motion and additive character evolution models are quite similar, they are computationally distinct, as Brownian motion is reliant on dated trees with informative branch lengths, and additive models only estimate intermediate states at nodes and are generally applied to undated trees (Farris, 1970; Hunt, 2007). This similarity thus suggests that modeling continuous characters under Brownian motion would likely be more suitable in a Bayesian framework, wherein the estimation of branch lengths is essential to the estimation of the tree.

The evolution of continuous variables under comparative methods has primarily been modeled under Brownian motion (i.e., random-walk; Felsenstein, 1973, 1985a; Gingerich, 1993) and Ornstein-Uhlenbeck (i.e., directional evolution; Beaulieu et al., 2012; Hansen, 1997; Butler and King, 2004), with many variants existing to relax or constrain model assumptions. These models allow for changes to accumulate continuously along branches, and because Brownian motion character evolution models conceptually mirror the additive method, character evolution models should be of similar utility in Bayesian inference as in parsimony. However, these analyses are typically post-hoc, and assume a tree known without error. Few have explored how these methods might behave in Bayesian or likelihood estimation of the tree using either empirical or simulation study. Parins-Fukuchi (2018) used a Brownian motion (Felsenstein, 1973) character evolution model in RevBayes (Höhna et al., 2016) to estimate trees from simulated morphological characters, demonstrating their utility in this type of analysis. Further work by Zhang et al. (2024) expanded upon this work with more general frameworks for adding continuous characters to trees. Brownian motion generally assumes that a trait will drift from its origin value with rate *σ*. This is a lightweight model, adding only one parameter, though it can be parameterized such that each character can have its own *σ* or *σ* can vary across branches on the tree.

Here, we explore the utility of using character evolution models of Brownian motion in estimating tip dated trees in a Bayesian framework under a Fossilized Birth-Death (FBD) model (Fig. 1). Our dated analysis can be readily separated into its component models: the substitution, clock, and tree models (Warnock and Wright, 2020). To jointly estimate the topology and node ages using the FBD, researchers generally assume a tripartite model of evolution: the first that describes the accumulation of character data, the second that describes the evolutionary rates, and the third that describes the speciation, extinction, and sampling events that generate the tree. The benefit of this type of joint model is that the researchers can treat each of the components as a discrete inferential module and also provides the flexibility to combine models according to the data in use (though see May and Rothfels (2023) for complications). This type of approach also allows for inference based on multiple types of data, including molecular or morphological character information. The model components are derived directly from the data, which can be multifarious and for our purposes here will be focused on morphological characters and fossil age information. In our case, this tripartite framework is a particular benefit because we can hold the clock model and tree model constant, while varying the input data and character models to investigate how combinations of data and model assumptions impact the resultant topology.

**Figure 1:**
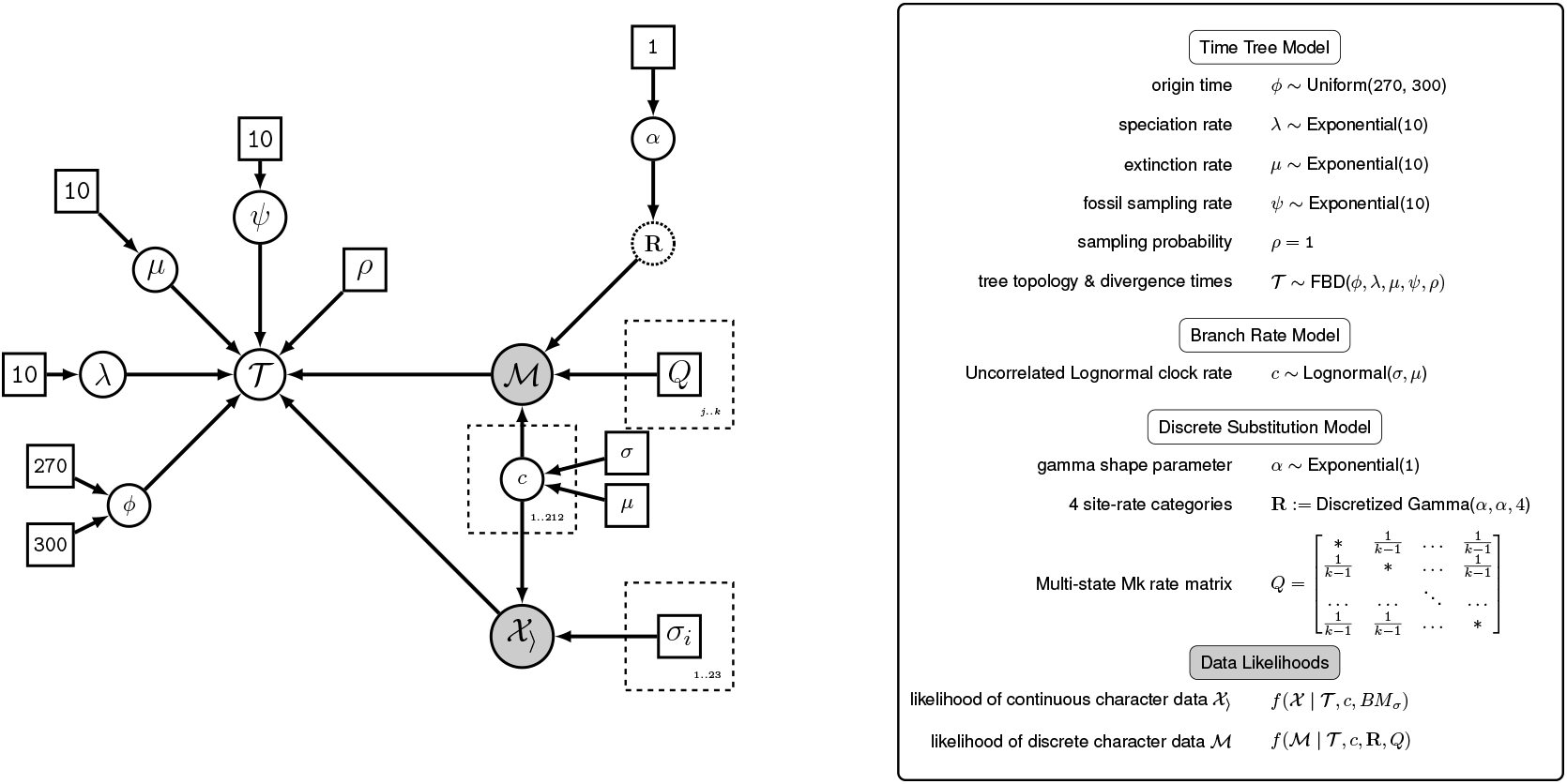
A graphical model displaying the components of the full joint phylogenetic model. An explanation of the model components is displayed at right. Adapted from Barido-Sottani et al. (2020).

### 1.1 Dicynodontia as a Study System

For these analyses, we rely on a recent phylogeny of Dicynodontia, a group of early synapsids (the mammalian total group) originating during the Permian Period. Dicynodonts were a diverse clade of herbivores known from the middle Permian through the Late Triassic periods (c.*∼*267 - 201.5 Ma), and can be readily recognized by their beaked jaws, usually without teeth other than the prominent maxillary tusks, efficient palinal feeding apparatus, wide skulls, and robust bodies (Angielczyk et al., 2017; Sulej and Niedźwiedzki, 2019). Over the past 30 years, the phylogenetic history of dicynodonts has been systematically untangled, with intensive alpha taxonomic revision that has reduced the overall species richness of the clade, but increased their utility for Permo-Triassic biostratigraphy (Kammerer et al., 2011; Wynd et al., 2017; Viglietti et al., 2016). Even following the extensive synonymization of historical taxa, however, dicynodonts remain one of the most speciose Permo-Triassic tetrapod groups, with over 100 valid species known globally (Fröbisch, 2009). The majority of dicynodont species richness occurs in two major radiations, the Permian therochelonians and their descendants, the Triassic kannemeyeriiforms. Permian and Early Triassic dicynodonts have an excellent fossil record, related to the exceptional nature of the Karoo Basin of South Africa preserving near-continuous sedimentation and capture of associated faunas during this interval (Smith et al., 2020b). Most Permian dicynodonts are known minimally from complete skulls, many are known from articulated skeletons, and some, such as *Diictodon* and *Lystrosaurus*, have sample sizes that are remarkable for a pre-Cenozoic terrestrial vertebrate (in the thousands of individuals) (Sullivan et al., 2003; Smith et al., 2013).

Dicynodontia represents an ideal test case to evaluate the impact of continuous characters on morphological character matrices, as they are completely extinct (necessitating the use of morphological data in phylogeny reconstruction), have a well-documented fossil record, and both continuous traits and discrete traits have been collected for this group and used in previous estimation of dicynodont phylogeny (Angielczyk and Kurkin, 2003; Kammerer et al., 2011, 2013, 2019). Current dicynodont phylogeny is the cumulative product of untangling over a century of taxonomic overprinting. The first iteration of the dicynodont data set analyzed herein was produced as part of a monograph revising the genus *Dicynodon*, which is known from hundreds of specimens from the late Permian previously attributed to more than 30 different species (Kammerer et al., 2011). Prior to the reassessment of *Dicynodon*, analyses of dicynodont phylogeny largely focused on either Permian or Triassic taxa in isolation, with examples covering both periods either incorporating very few taxa from the non-focal period or generating composite trees from separate analyses (for a deeper discussion on the origins of this tree, see the section “Phylogenetic Analysis” in Kammerer et al. (2011) and the associated citations therein).

All analyses of dicynodont phylogeny from the past decade are based on the underlying dataset of Kammerer et al. (2011), which introduced continuous character data to this system. The goal of the Kammerer et al. (2011) data set was to broadly sample dicynodont diversity, providing for the first time nearly comprehensive coverage of the clade throughout its Permian and Triassic history. Making up this data set were a combination of novel characters and characters adapted from previous analyses of dicynodont phylogeny (e.g., Maisch (2002); Damiani et al. (2007)), encompassing the wide variety of morphologies observed throughout anomodont evolution. However, due to variation in the quality of the dicynodont fossil record over time, different character types vary in importance across different groups of dicynodonts.

Continuous variables were introduced to the data set to address historical inconsistencies in discretizing numerous features better explained as measurements and ratios than discrete-state characters. When incorporated under an additive model alongside morphological character data in parsimony-based phylogenetics, these characters have been demonstrated to provide additional phylogenetic resolution unavailable using only discrete-state characters (Kammerer et al., 2011). This is especially true for taxa with very incomplete codings for the majority-cranial discrete-state character set. The later Triassic kannemeyeriiforms (particularly the Stahleckeriidae) tend to be known from less complete crania than their Permian predecessors, with greater representation of taxa based on fragmentary postcrania instead of complete skulls or articulated skeletons. Some of these taxa are currently represented only by single damaged skeletal elements (e.g., *Eubrachiosaurus browni*) or extensive but eroded and disarticulated remains (e.g., *Placerias hesternus*) (Kammerer et al., 2013; Fiorillo et al., 2000). Observed differences in postcranial anatomy between these taxa are largely proportional, and continuous character measurements (primarily ratios) have been an important component of character support in this part of the tree. This offers a unique opportunity to explore which group of characters is contributing signal to the estimation process and how conceptually similar models results differ from one another. Not only will this work help establish future models in phylogenetic estimation, but also we have the unique opportunity to question our established methods and whether or not they behave the ways we expect.

## 2 Materials and Methods

### 2.1 Character Evolution Models

#### 2.1.1 Discrete Characters: Mk Model

Discrete characters are generally modeled using the Mk model in a likelihood-based framework. The Mk model (Lewis, 2001) is a Markov model for morphological evolution, serving as a morphological analogue to the Jukes-Cantor Model (Jukes and Cantor, 1969). Due to the inherent complexity and diversity of morphological evolution across different characters, the assumptions of the Mk model are inherently simplistic. Notably, this model assumes that the rate of change between the character states are equal.

In the Mk model, the *k* denotes the number of possible states for a given character, allowing the model to be flexible and accommodate varying numbers of character states across different characters. The rate matrix (Q-matrix) for this model can be generalized based on the number of character states, as follows:

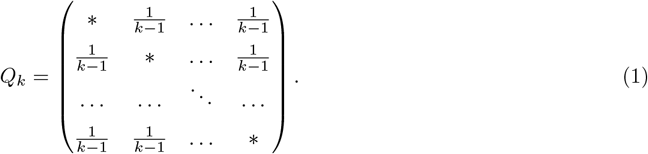

To address the biological oversimplifications inherent in the Mk model, various extensions have been proposed. For instance, Nylander et al. (2004) introduced priors on the stationary frequencies to relax the assumption of equal transition rates, but were not used in this paper, as prior work demonstrates that character model misspecification is largely unproblematic in dated analyses (Klopfstein et al., 2019).

We employed the Mk (equal-rates) model to generate the substitution (Q) matrices for each character. Partitions were made that separate each character based on the number of character states (Khakurel et al., 2024; Mulvey et al., 2024), to establish *n* Q-matrices, where *n* is the maximum number of states for any character in the dataset. For the Mk model, we established among-character rate variation by using gamma distributed rate variation with four discrete rate categories (Yang, 1994).

#### 2.1.2 Brownian Motion

The Brownian motion model (Felsenstein, 1973), also known as the random walk model, assumes that trait evolution occurs through random, incremental changes over time, with both the direction and magnitude of each change governed by a stochastic process. This model includes a single parameter, *σ*, which represents the evolutionary rate and dictates the size of the phenotypic shift in each time step. Under this model, the phenotypic change in a descendant lineage is assumed to follow a normal distribution, with the mean centered on the ancestral state and the variation proportional to the product of *σ* and branch length. Consequently, traits are expected to exhibit greater changes along longer branches, reflecting the increased time elapsed since divergence from a common ancestor. This model has been used inferentially in prior studies (Parins-Fukuchi, 2018; Zhang et al., 2024).

As a model driven purely by drift, the Brownian motion model assumes that the trait evolution is stochastic, independent of other factors such as natural selection or genetic constraints (Butler and King, 2004). Despite this simplification, the model remains a valuable tool for investigating general patterns and trends in the evolution of continuous traits over extended time scales.

We used two different types of Brownian motion model:

- One-Rate Brownian Motion: Under this model, all characters share a *σ* value drawn from a uniform distribution bounded between 1e-5 and 1e-1. This model will be referred to as **One-Rate** in figures and further text.
- Multi-Rate Brownian Motion: Under this model, each character has an independent *σ*, drawn from the uniform distribution described above. This model will be referred to as **Multi-Rate** in figures and further text.

Both models assume uncorrelated characters. This will strike some readers as unrealistic, but for a first pass application, this seems reasonable, and matched to the assumptions of independence we make for discrete characters.

#### 2.1.3 Fossilized Birth-Death Model

The FBD model was held constant across all datasets and analyses. For the major parameters of the FBD, we used the Paleobiology Database (PBDB) visual tools (Peters and McClennen, 2016; Uhen et al., 2023) to obtain estimates for origination and extinction rates for the clade from published specimens. We sampled speciation rates from an exponential prior with a mean of 5.5, based on this output. Turnover rate was pulled from a uniform distribution between 0.90 and 1.05. From turnover rate and speciation rate, we can estimate extinction rate (turnover - speciation rate) as well as diversification rate (speciation - extinction rate). The fossil sampling rate (*ψ*) was sampled from an exponential distribution with a mean of (1/0.072), which is the average per-million year expectation given the character frequencies. We set the proportional taxon sampling of the youngest time slice (*ρ*) at 0.231, based on the distribution of finds among synapsids and archosaurs from sediments contemporaneous with the latest therapsids. Origin time, the time of divergence for the last universal common ancestor for the given topology, was sampled between 270 and 300 million years old. A complete model can be viewed on Fig. 1.

No clade constraints were used in our analyses when comparing the continuous, discrete, and joint models to one another. Rather, starting trees where drawn from a distribution of trees plausible under the FBD distribution. We ran the analysis with and without sampled ancestors. For the fossil ages, we used occurrences from the Paleobiology Database (PBDB), including many that were entered for the purpose of this project. Because the PBDB returns only general ages in its output, we used a separate database (see Congreve et al., 2021) that includes biozonation, radiometric dates (primarily U/Pb), and sequence stratigraphy. The estimated first-appearance age for each taxon is pulled from a uniform distribution, where the ranges are the estimated time for the most recent common ancestor for the given clade (TMRCA in `RevBayes`), minus the maximum and minimum ages. This standardizes the age of each taxon based on the clades they belong to.

One possible concern is that some of the taxa analyzed here are themselves zone taxa. For example, the *Dicynodon* zone (= *Dicynodon-Theriognathus* Subzone of the *Daptocephalus* Assemblage Zone) is thought to span from 255.2 - 254.0 Ma (Smith et al. 2020a). Simply assuming that all *Dicynodon* occurrences fall within that span would present circular reasoning. However, the zone taxa analyzed here do not have a 1:1 relationship with their eponymous zones; the biozones in question represent characteristic assemblages (presence of multiple taxa simultaneously, some components of which pre- or postdate the nominal zone). Other criteria used to date PBDB sites (including U/Pb data) are sufficient to avoid possible circularity.

All the original data and scripts necessary to reproduce the analyses reported in this study can be accessed through the Dryad link: http://datadryad.org/stash/share/ahyNsjmSGsFvdaQ0SvVhmUj2Jmp0hnhnlK8Wlp_ensQ.

#### 2.1.4 Clock Models

We used an uncorrelated relaxed lognormal clock to model the rate of evolution across the tree (Thorne et al., 1998; Drummond et al., 2006). This clock model is very flexible, allowing ancestor and descendant branches to have very different rates of evolution. This model was held constant across all analyses and datasets.

#### 2.1.5 Joint Inference

We ran a total of five different models that include varying amounts of discrete and continuous character information (Table 1): continuous with all characters sharing a Brownian motion rate (**Continuous - One-Rate**), continuous with characters having their own BM rate (**Continuous - Multi-Rate**), discrete characters under the Mk model and continuous characters in which all continuous characters share a BM rate (**Combined - One-Rate**), and discrete and continuous characters with continuous characters having their own BM rate (**Combined - Multi-Rate**), all of which were compared to the discrete-only (**Discrete**) model.

**Table 1:** A table demonstrating the various character models tested in the paper.

| Name | Discrete Characters Included | Continuous Characters Included | Number of $\sigma$ Brownian Motion Rates | Generations to Converge |
| --- | --- | --- | --- | --- |
| Continuous - Multi-Rate | Yes | No | 23 | 135,000 |
| Continuous - One-Rate | Yes | No | 1 | 235,000 |
| Combined - One-Rate | Yes | Yes | 1 | 200,000 |
| Combined - Multi-Rate | Yes | Yes | 23 | 135,000 |
| Discrete | Yes | No | NA | 260,000 |

We performed phylogenetic inferences in RevBayes (Höhna, 2014; Höhna et al., 2016), which allows for easily customizable scripting and extensive researcher control over parameterization. We use continuous time Markov Chain Monte Carlo (MCMC) to explore tree space (see Wright (2019) for more on the MCMC and its utility in morphological phylogenetics and Barido-Sottani et al. (2024) for a general overview of MCMC in phylogenetics). A key aspect of the MCMC is that no two analyses will be identical but when modeling the same phenomenon one expects different analyses to converge to the same posterior distribution. Therefore, we ran two replicates of each analysis, and checked for convergence both within and between replicates. To evaluate whether or not our model has reached convergence, we use estimated sample size (ESS) as an initial marker, with an ESS of 200 being considered the threshold minimum for convergence (Lanfear et al., 2016; Rambaut et al., 2018). We checked convergence of the tree itself in `RWTY` (Lanfear et al., 2016).

We generated summary maximum clade credibility trees for each analysis using the `mccTree` command in `RevBayes`. To do this, we discarded 25% of the sample as burn-in and combined both replicates of each analysis.

### 2.2 Post-Analysis Processing

Following our estimations, we quantified several summary statistics from each run. In RevGadgets, we calculated tree and branch lengths for all trees in the posterior distribution (Tribble et al., 2022). We also used the software RWTY (Warren et al., 2016) to calculate a treespace visualization (Hillis et al., 2005; Whidden and Matsen IV, 2015). For the treespace visualization, we recomputed our MCMC tree logs with sampled ancestor printing suppressed (but sampled ancestors still allowed in the analysis). This is because few software packages accept sampled ancestors. We sampled the chains every 100 steps, and calculated Robinson-Foulds distance (Robinson and Foulds, 1981) both within each analysis type (i.e, Combined-Multirate) and between analysis types. We then plotted these to treespace using multi-dimensional scaling to plot 100 points per analysis type to a 2-D space (Warren et al., 2016). The resultant graphic was refined using ggplot2 (Wickham, 2011) for custom colors.

## 3 Results

### 3.1 Model results

All model and data combinations ultimately reach convergence in our experiments, as can be seen in Fig. S1. Plots on Fig. S1 are shown downsampled, as to appear on the same axis. However, in Table 1, the number of generations required for convergence is shown. Interestingly, the discrete-only datasets take the longest to converge, followed by the combined - one-rate analysis. The combined - multirate analysis takes the least time. It should be noted that generations in `RevBayes` run multiple moves per iteration (Höhna et al., 2014). Thus, these are not comparable to other software packages that can use continuous traits, such as `BEAST 2` (Bouckaert et al., 2014).

As shown on Fig. 2a, the different treatments arrive at different origin times for the clade. Most treatments favor an origin time of between 280-270 million years before the present, which is consistent with the estimated age of divergence between the major therapsid subclades (Biarmosuchia, Dinocephalia, Anomodontia, and Theriodontia) based on recent analyses examining therapsid interrelationships more broadly (Matamales- Andreu et al., 2024). The discrete and combined-multirate datasets and models prefers a slightly younger origin time, at around 270 Ma. These ages accord with prior work, as the oldest fossil anomodont (the larger clade containing dicynodonts and their nearest relatives) in our sample, *Biseridens qilianicus*, has an estimated age of only *∼*265 Ma and is morphologically very similar to basal members of other therapsid subclades (particularly dinocephalians), suggesting that there was not a protracted period of divergence prior to its appearance in the record. Even with differences in the origin times preferred by different models, there is substantial overlap between the youngest estimates and older ones.

**Figure 2:**
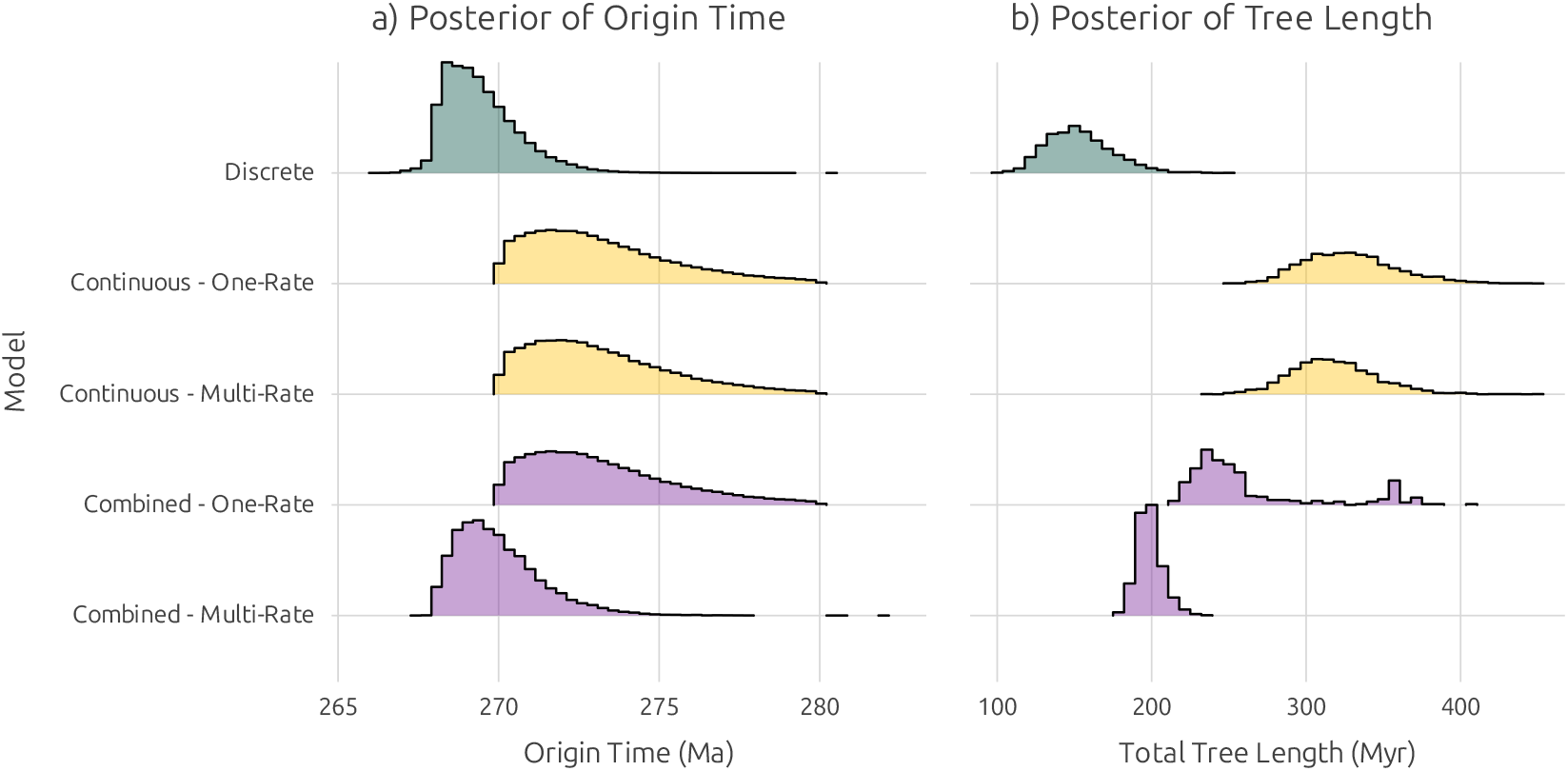
Visualization of posterior distributions of a) origin times and b) tree lengths (in millions of years before the present) between different models and data sources.

Although we can easily see the distribution of origin times, the same is not true for major groups of organisms without instituting clade constraints. Because each tree in the posterior may have slightly different arrangements of species, it is unlikely that a single node in the analysis consistently retains the same clade across the entire posterior. Therefore, we can only use the maximum clade credibility trees for our given treatments to evaluate how divergence dates correspond to present estimates. We checked the divergence dates across analyses for the following major clades in the taxon sample: Anomodontia, Dicynodontia, Bidentalia, Dicynodontoidea, and Kannemeyeriiformes (Table 2). The clade ages in the discrete model are largely consistent with the expected dates, albeit with Dicynodontia being younger than expected, with a rapid diversification from the Anomodontia one million years prior. Overall, the estimated ages are largely consistent with expected ages, with the most notable exceptions being the age for the Bidentalia, which was consistently recovered as older in all but the discrete analysis. This discrepancy is largely the product of members of the bidentalia being recovered among early diverging anomodonts in the continuous and combined analyses, which artificially pulls these clades closer to the age of divergence for the Dicynodontia. Outside of the age estimate for the Dicynodontoidea (specifically the continuous-multi rate model), ages are within 2-3 million years of expected divergence dates, which is a reasonable error given the inherent error expected from U/Pb radiometric dates for late Paleozoic and early Mesozoic deposits. The topologies for the continuous (both) and the combined multirate analyses dissolve many of these major clades, and so ages are more reflective of earliest clade member ages, rather than the estimated age of diversification of a monophyletic lineage.

**Table 2:** Age estimates of maximum clade credibility trees across models compared to expected ages.

| Clade | Expected age (Ma) | Discrete age (Ma) | Continuous single rate (Ma) | Continuous multirate (Ma) | Combined single rate (Ma) | Combined multirate (Ma) |
| --- | --- | --- | --- | --- | --- | --- |
| Anomodontia | 274 – 270<sup>*1</sup> | 272 | 275 | 273.5 | 269.5 | 273 |
| Dicynodontia | 269 – 265<sup>*2</sup> | 271 | 272.5 | 271 | 268.5 | 272 |
| Bidentalia | 263<sup>*3</sup> | 262 | 268.5 | 269.5 | 266.5 | 269 |
| Dicynodontoidea | 261<sup>*4</sup> | 259.5 | 263 | 266 | 259 | 261 |
| Kannemeyeriiformes | 250<sup>*5</sup> | 249 | 252 | 251 | 248 | 250 |

Age estimates are based on divergence dates approximated from trees depicted within the supplemental materials. Age estimates are based on figures or text in <sup>\*1</sup>Matamalas-Andreu et al. (2024), <sup>\*2</sup>, <sup>\*3-4</sup> Angielczyk et al. (2021), <sup>\*5</sup> Sulej and Niedźwiedzki (2019). Ages are approximations and should be viewed as having an expected error of $\pm 1$ Ma

### 3.2 Tree topology

Our analyses did not reproduce the consensus tree that has been produced by previous cladistic analyses and consistently supported by years of expert opinion (Olivier et al., 2019; Angielczyk and Kurkin, 2003; Angielczyk et al., 2014; Kammerer et al., 2011; Martinelli et al., 2021). We find that, in general, when continuous variables are examined alone, the produced trees are essentially nonsense, dissolving expected clades and yielding taxon groupings with minimal morphological similarity and high levels of stratigraphic debt (Fig. 3; for fully labeled trees see Fig. S2 - Fig. S6). This is best exemplified in the Kannemeyeriiformes; this clade is known only from the Triassic Period and its members have consistently been recognized as more closely related to one another than to Permian dicynodonts (e.g., Kammerer and Ordoñez (2021)). When only continuous data are incorporated, the Kannemeyeriiformes are recovered as a polyphyletic assemblage nested in two independent lineages of Permian dicynodonts. When the discrete data are examined alone, we find that Kannemeyeriiformes remains mostly intact, but a few kannemeyeriiform taxa are instead reconstructed as sister taxa to Permian dicynodonts traditionally recovered as distant to this group. Generally, however, the results of this analysis are far more similar to established morphological and stratigraphic relationships for dicynodonts than the hypotheses presented in the continuous-only data. Independently, the impact of variation in fossil record quality is evident in the characters, where expected relationships amongst Permian dicynodonts are more or less conserved under the discrete characters, and the continuous variables assist most strongly in determining relationships among the Triassic dicynodonts.

**Figure 3:**
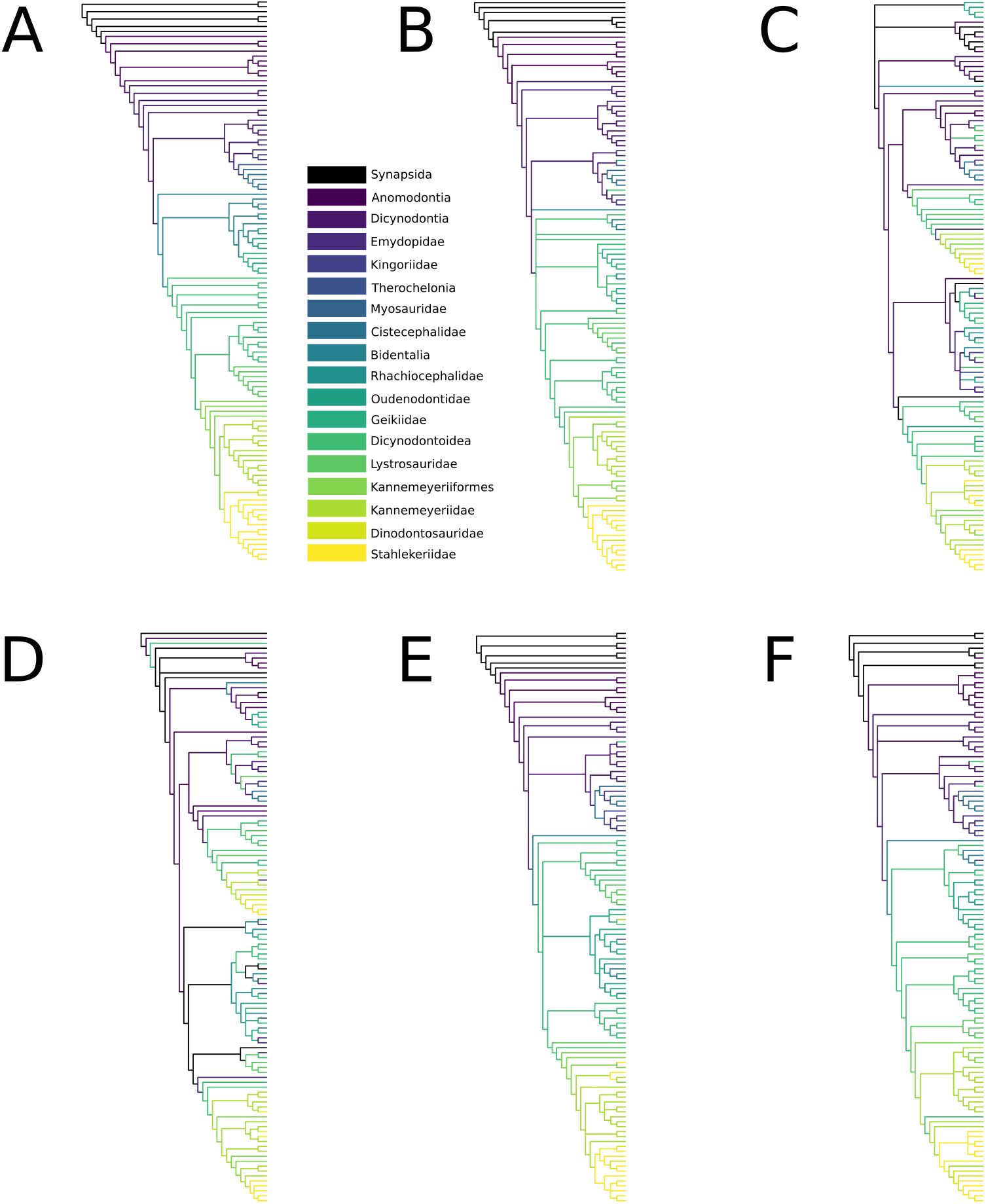
Topological differences between models plot depicting how dicynodont groupings are differently dissolved in our various analyses. A) current consensus topology. B-F are maximum clade credibility trees (MCC) from our five models: B) Discrete only; C) Continuous One-Rate; D) Continuous Multirate; E) Combined One-Rate; F) Combined Multirate. The currently accepted topology was drawn based on Kammerer et al. (2016, 2019); Kammerer and Ordoñez (2021). Topologies B-F were redrawn from our MCC trees. Full trees with tip annotations and clade supports are included as S2 - S6. Tips for the groupings “Synapsida”, “Anomodontia”, “Dicynodontia”, “Therochelonia”, “Bidentalia”, “Dicynodontoidea”, and “Kannemeyeriiformes” represent the basal grades within these groups exclusive of the other listed clades.

However, even the discrete data yield problematic taxonomic relationships at the base of Anomodontia and the dicynodont subclade Emydopoidea, specifically concerning the positions of the genera *Ulemica* and *Dicynodontoides*, respectively. *Ulemica* is a non-dicynodont anomodont, historically recovered in the clade Venyukovioidea; indeed, the primary basis for this clade’s existence, as the well-preserved, extensively studied remains of this taxon were the basis for allying taxa based on more fragmentary remains such as *Venyukovia* and *Otsheria* (Ivakhnenko, 1996). It should be noted that the posterior support for *Ulemica* is consistently low (*<* 0.04), suggesting that its placement is spurious, and should be taken lightly.

Some differences in topology relative to the ‘standard’ parsimony-based trees are expected for our analyses; for example, while the recovered position of *Niassodon* is also problematic, this has also been a labile taxon in previous parsimony analyses of dicynodonts. Similarly, the failure to recover a monophyletic Cryptodontia is an issue that has also arisen in various prior iterations of dicynodont phylogeny (Angielczyk et al., 2017) and should not cause major concern here. By contrast, the ‘erroneous’ positions of exceptionally well-represented taxa such as *Ulemica* and *Dicynodontoides*, with extremely well-supported placements in previous trees, is highly concerning, and suggests that the explanatory power of synapomorphies in parsimony analyses do not readily transfer to Bayesian total evidence analyses.

### 3.3 Tree lengths

To gain a sense of how model choice impacts phylogenetic inference, we estimate tree length as a cumulative sum of all internal branches within the tree. The discrete produces the youngest set of trees. Continuous characters (only) modeled under either the one-rate or multirate models favor longer trees. Combined analyses split the difference, with multirate models being closer in length to discrete trees. This is consistent with our findings on origin time (Fig. 2a), which reflect the continuous-only datasets favoring an older age. Overall, including continuous and discrete traits together seems to moderate the conflicting signals of age from the two trait types on their own.

As discussed in section 2.1.3, we ran the analyses with and without sampled ancestors. We decided not to visualize them in our tree figures as they were numerous and poorly-supported. On Fig. S7, we plot the ages of the MCMC runs with and without sampled ancestors. We find that without sampled ancestors incorporated in the model, trees grow unrealistically long. This is largely due to inflating the number of bifurcations, and therefore pushing back divergences deeper in the tree.

### 3.4 Treespace

When we generate a treespace (Fig. 4) comparing all five experimental treatments, we see a pattern not dissimilar from that seen on Figs. 2a, and 2b. The combined multirate and one-rate cluster together, topologically, with the discrete characters. The continuous data types are further away in space, indicating less similar trees to the combined and discrete cluster. However, the continuous trait trees are still quite dissimilar from one another, as evidenced from their wide dispersal.

**Figure 4:**
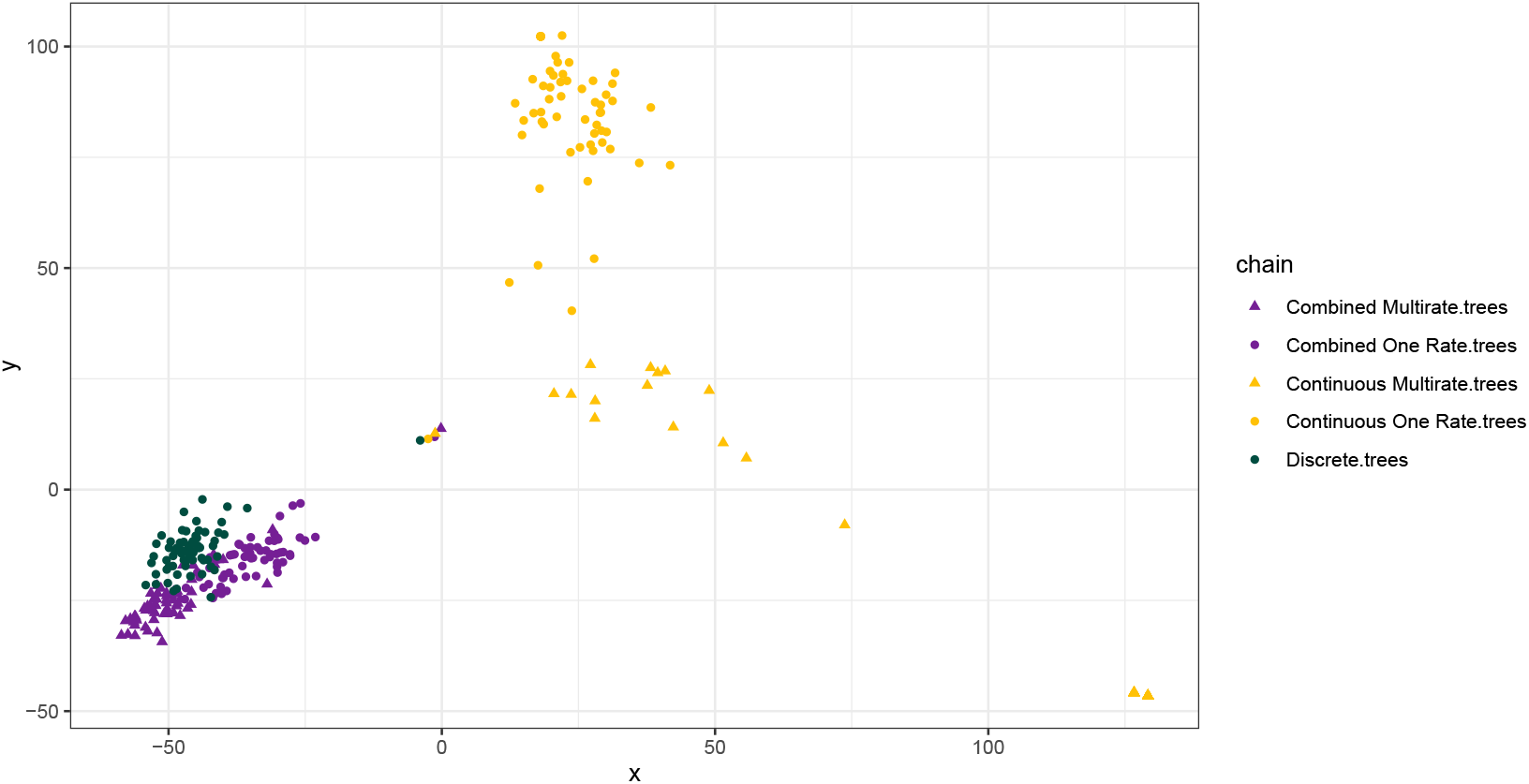
A treespace depicting how the five different experimental treatments sampled tree space. Points are colored by treatment. The metric is Robinson Foulds, which is the topological difference between two trees weighted by branch length similarity. Thus, points that are closer together are more similar topologically and in branch lengths.

## 4 Discussion

### 4.1 The unintended consequences of continuous characters

Continuous variables preserve morphological characteristics via direct measurement. It would follow that such quantification would have utility in morphological phylogenetics, but we find the answer to not be so clear-cut. Inclusion of continuous characters appear to have a large effect on topological results and the manner in which treespace is explored (Fig. 4). We see this in the dissolution of clades, such as Geikiidae and Lystrosauridae, being interspersed throughout the tree when continuous variables are included (Fig. 3, for full, labeled trees, see Figs. S2 - S6). Geikiids are Permian dicynodonts that are short and broad-skulled, and in their slightly less inclusive subclade, Geikiinae, these taxa possess cranial ornamentation in the form of nasal and parietal bosses (Kammerer and Angielczyk, 2009). The dissolution of geikiines is thus problematic because it would require numerous independent events wherein major morphological modifications to the nasal bones would need to occur; ultimately the present rarity of this characteristic in non-geikiine dicynodonts suggests homology of the nasal bosses as a better supported hypothesis. The

Lystrosauridae is clade of dicynodonts that spans the Permo-Triassic Mass Extinction event and are often regarded as globally cosmopolitan, generalist disaster taxa in earliest Triassic sediments (Viglietti et al., 2021). Although the exact position of the Lystrosauridae in greater dicynodontoid phylogeny has been poorly supported in previous analyses, the monophyly of the Lystrosauridae has been empirically supported repeatedly (Maisch, 2002; Kammerer and Angielczyk, 2009). These clades are well-supported in the 174 characters of the discrete-only analysis, but it only takes the addition of the 23 continuous characters to break that support. This finding is reminiscent of similar findings in combined molecular-morphological datasets (Bapst et al., 2018), in which a few morphological characters can break support for clades supported by larger volumes of molecular data. This is particularly problematic, as the continuous variables are largely used to better place the kannemeyeriiform dicynodonts and their generally fragmentary preservation when compared to Permian dicynodonts. Although there are species of kannemeyeriiform dicynodonts known from complete skeletons (e.g., *Shaanbeikannemeyeria xilougouensis*, (Liu, 2022); *Ishigualastia jenseni*, (Cox, 1965); *Kannemeyeria simocephalus* (Govender et al., 2008)), much of the kannemeyeriiform record is comprised of incomplete skeletons and often fragmentary elements (e.g., *Argodicynodon boreni* (Mueller et al., 2023); *Eubrachiosaurus browni* (Kammerer et al., 2013); *Sungeodon kinkraemerae* (Maisch and Matzke, 2014)). Given this, diagnostic cranial features are missing in some taxa, whereas measurements and ratios can still be coded for their fragmentary postcrania. Thus, with discrete characters underrepresented in the kannemeyeriiform data, these continuous characters are expected to have a proportionally larger impact on kannemeyeriiform relationships than they would on non-kannemeyeriiform dicynodont relationships. This group is non-monophyletic when exploring the discrete-only model, and are broken up more significantly throughout the tree when continuous traits are included. The fact that these characters essentially produce a result opposite to their intent when treating them as character evolution models should raise caution concerning the use of such data in any phylogenetic context.

The complications surrounding continuous variables in phylogenetics, especially in the context of branch lengths, is potentially due to model misspecification. When examining our trees, we find what appear to be opposing results when continuous characters are treated under a single rate or individual rates for each continuous character, particularly when in combination with discrete characters (Fig. 3). This suggests that a single-rate Brownian motion model is likely to be inappropriate for all characters. This is perhaps unsurprising, as we do not assume all discrete characters evolve at the same rate (Yang, 1994; Wright and Wynd, 2024) We also see this impact in the parameters of the FBD model - when we use a multirate model for combined discrete and continuous datasets, the origin time aligns very well with the known fossil record for the group (Fig. 2a; Fröbisch, 2009). This largely also accords with the discrete-only age.

However, when only a single rate is imposed, the model struggles to find reasonable topologies that satisfy each and every continuous variable sharing the same history. When we incorporate discrete characters, it is simpler to include the continuous characters with a one-rate BM. The model has less difficulty finding starting parameters that are computable. This is likely due to the simplicity of the model space as a result of only adding one parameter. When the multirate BM model is implemented, 23 separate parameters need to be adjusted each iteration, increasing model complexity and computation time. However, despite this complexity, multirate model reaches convergence in far fewer generations once it begins running (Table 1). As we will discuss below in the section **4.3**, the multirate models produce more reasonable estimates along several fronts, and we recommend exploring them in analyses incorporating continuous traits, a finding that has recently also been reported in Wright and Hopkins (2025), in which the authors corroborate out finding that continuous and discrete analyses often search treespace in very different ways.

Character evolution models are extremely powerful tools when the model of evolution properly matches the history of the trait. Existing character evolution models include those that explore random walks of evolution (i.e., Brownian motion), where the inherent assumption is that this is a trait that evolved largely via genetic drift. This can be a serious issue if the evolutionary history of the trait is non-Brownian, or if the evolution is directional (i.e., Ornstein-Uhlenbeck character evolution models (Hansen, 1997; Butler and King, 2004)). This kind of character model is more appropriate when applied to a trait that appears to have an optimum (*θ*). This is not to say that there is a true optimum, but that there was a selective pressure on that trait, which eventually reached a point of general stability. The OU model assumes rapid evolution early in the trait history followed by relatively slow change once the optimum is reached. Brownian motion is a memory-rich process where the current state is dependent on the state prior to it. With stochastic change centered around a mean of 0, the model depends on small differences or long branches to accumulate changes, such that the root state of the tree is informative for the tip state of the traits. OU models, however, are memory rich in the stability phase, but large shifts during the directional evolution phase suggest that the root state is less useful when estimating characters that do not evolve by random evolutionary processes. OU processes may be a better way to model continuous traits for phylogenetic inference, but are not currently implemented for inference to test.

Our analyses with continuous traits still produce interesting artifacts. As seen on Fig. 2b, continuous traits run under either multirate or one-rate Brownian motion processes produce much longer tree lengths than either discrete or combined discrete-continuous datasets. Conceptually, this makes sense as to why it would be a problem in the Bayesian analysis, but not the cladistic analysis *sensu* Farris (1970). Because the additive method only requires tips, and nodes, the differences between the tips and nodes would be specific to each individual character, such that characters with vastly distinct values and rates of change can be independently estimated using the same algorithm. Thus, the additive method of Farris has more in common with the multirate than one-rate BM. In the case of strict BM, the focus on branch lengths means that a single rate parameter, is unlikely to properly explain the evolutionary history of completely different traits, let alone, 23 different traits encompassing different skeletal elements.

Potentially the most impactful biases of the fossil record is preservation (Webster and Hughes, 1999). While we have thus far extolled the objectivity of continuous trait data, they are not without biases. For both the vertebrate and invertebrate fossil records, specimens are rarely preserved as they would have been in life. Whether it be soft tissues that are inherently unlikely to preserve, broken or partially preserved elements, entirely missing elements, or distortions from shifts to the local geography, fossilization obliterates and obscures the biological entities it captures. Although continuous trait measures may have less of a human interpretive filter, raw measurements taken directly from fossil specimens are rarely to be trusted without some degree of scrutiny. Common approaches to accounting for taphonomy—the process of fossilization— include retrodeformation analyses (Arbour and Currie, 2012; Kammerer et al., 2020; Angielczyk and Sheets, 2007; Schlager et al., 2018; Lautenschlager, 2016), extrapolations based on numerous specimens (Klingenberg, 1996; Klingenberg et al., 2001; Lande, 1979; Alexander, 1985), and our own work using linear mixed effects models to account for random deformation (Wynd et al., 2021). These methods are potentially of use to researchers looking to use continuous traits inferentially.

However, each of these approaches carries their own pitfalls that limit broad application for small datasets, which account for the vast majority of existing paleontological datasets (Koch, 1991). Because raw measurements of fossils still include significant bias, phylogenetic characters have been primarily based on observations that can be discretized (Baum, 1988). It is clear that discretized characters will always be a staple of phylogenetic analyses based on fossils; regardless, the incorporation of continuous variables is critical to maximize the amount of data that can be brought to bear on a problem.

### 4.2 The benefits of including continuous variables in joint phylogenetic estimation

The issues pointed out above bear on the signal within continuous characters and how we need to thoroughly vet both our data and how it behaves under differing model circumstances. As seen on (Figs. 2a and b), continuous traits can have a powerful impact on the parameters of our models, and on the way treespace explored (Fig. 4). If continuous traits had no signal, we would not observe these differences. Therefore, it is crucial to think carefully about the choices used to model them. When only continuous characters are considered, trees are much longer (Fig. 2b) than when datasets are combined or discrete. Importantly, when more than a single rate is specified for the Brownian model in a combined dataset, there is no major change to the origin time, and branch lengths are consistent with those generated through discrete characters under an Mk model (Figs. 2a and b).

We discuss above how, aside from branch lengths, the additive model is conceptually similar to a character evolution model following multirate Brownian motion. However, the additive model as a parsimony-based method lacks a unifying rate parameter, thus the additive model can effectively incorporate characters with lots of rate heterogeneity. The BM character evolution model, on the hand, requires some value to ascribe to the rate of change for the trait in question; and when only a single rate is given for over 20 characters, many of which sample completely different skeletal elements, it is unlikely that those characters are going to contribute meaningful information, and will instead produce erroneous clades. This may seem discouraging, but the opposite is actually true. It should be very validating that the model fails so catastrophically when we try to shoehorn fundamentally distinct data into the same parameters and expect the model to appropriately sample the nuance. This might be considered similarly to not allowing among-character rate variation in a substitution model (Yang, 1994; Capobianco and Höhna, 2025; Wright and Wynd, 2024). The addition of discrete traits as a separate partition substantially helps inference in both the one- and multi-rate models.

### 4.3 Insights based on dicynodont evolution

When it comes to dicynodont topology, the discrete and combined-strict analyses produce topologies most consistent with the currently-supported dicynodont relationships. For the majority of Permian dicynodonts, the historically recognized clades are largely preserved and the species based on more complete specimens tend to be placed in their expected positions. However the placement of some taxa should raise concerns. Specifically, the genera *Ulemica* and *Dicynodontoides* are some of the better sampled taxa in our dataset, and in parsimony analyses, their positions are supported by minimally dozens of synapomorphies, yet their phylogenetic positions in our analyses are highly variable and are supported by extremely low posterior probabilities. *Ulemica* is recovered well-nested within the Dicynodontia in the discrete-only analysis, but is pulled closer to its ‘proper’ position when continuous characters are incorporated; however, the continuous-only analyses create entirely unrealistic topologies in the rest of the tree. The recovery of an ‘erroneous’ (relative to established phylogenies) position for *Ulemica* is notable because it is known from a reasonably complete skull allowing it to be coded for the majority of characters in the analysis. Its position in parsimony-based analyses as a taxon falling outside of Dicynodontia is supported by 33 discrete dicynodont synapomorphies not present in *Ulemica* (based on analysis of Kammerer et al., 2019). Given the strong support in previous analyses, it is both unlikely and strange that *Ulemica* would behave as a rogue taxon with such low stability between analyses, especially compared to taxa known from much more fragmentary material. The case for *Dicynodontoides* should be even more concerning, as this taxon is one of the most complete taxa sampled in our analyses, being scored for over 95% of characters, including extensively studied postcranial skeletons (Angielczyk et al., 2009). Typically a non-bidentalian emydopoid, it is recovered here as belonging to Bidentalia (Discrete-only and Combined Multirate), and in some analyses more specifically among lystrosaurids (Continuous only One-Rate and Multirate) or as sister taxon to geikiids (Combined One-Rate). These latter two groups ae supported by numerous synapomorphies in previous analyses that are not present in *Dicynodontoides*, and in no prior analysis has it been recovered as closely related to them (pers. obs. CFK).

Given the completeness of these species, we lack any strong arguments as to why these taxa seem to have so much lability; it is possible that the explanatory power of synapomorphies in parsimony analysis does not appropriately transfer to Bayesian total-evidence analyses, but this will require more targeted studies to address at a broader scale. Outside of *Ulemica* and *Dicynodontoides*, the Bayesian discrete and combined analyses tend to largely support the expected clades, with most other rogue taxa having a known record of phylogenetic lability under parsimony analyses. Because the driving force behind the use of continuous characters in this data set is to better place the Kannemeyeriiformes, these results make intuitive sense. When discrete characters are included, in general, the species known from the most complete specimens are in or near their expected phylogenetic positions. With the continuous characters however, the impacts of model misspecification are directly represented on the phylogeny via the ‘incorrect’ placements of kannemeyeriiform taxa. Although both the strict and multirate continuous-only models produce trees that are effectively nonsense, the multirate model tends to preserve smaller clades and intersperse them throughout the tree in seemingly random clades. The strict model on the other hand, tends to take more individual species and intersperse those throughout the tree. We interpret this difference as the strict model attempting to ‘shoehorn’ characters that likely evolved at a different rates from one another into a single explanatory rate; this is an issue that is not present in the parsimony analysis because the additive character model of popular tree-building software TNT is based on minimizing the between node differences across the tree rather than attempting to apply a rate to a branch length.

In the divergence dates, the different analyses produce dates more or less in line with prior work. Each of the analyses produces an age for Anomodontia that are largely consistent with recent tip-dated and time-calibrated phylogenies (Angielczyk et al., 2021; Sulej and Niedźwiedzki, 2019; Matamales-Andreu et al., 2024). Except for the combined multirate analysis, the age of the Dicynodontia is consistently older than would be expected, and is largely driven by non-anomodontian therapsids being interspersed throughout early anomodonts and dicynodonts, artificially pulling the age of Dicynodontia back. The same can be seen in the Bidentalia, but only the discrete analysis is consistent with the expected age. This is largely driven by the dissolution of the Cryptodontia, the sister taxon to the Dicynodontoidea. Across analyses, major groups of Cryptodonts are largely retained (e.g., the Oudenodontidae, Rhachiocephalidae, and Geikiidae), but these clades are only joined in a common clade in the discrete analysis. Geikiids tend to be pulled into the Dicynodontoidea as their earliest diverging members, which would cause the divergence date for the clade to be overestimated. A similar pattern is seen in the Kannemeyeriiformes, where kannemeyeriids and stahleckeriids are not found as sister taxa, but are frequently nested within subsets of dicynodontoids, a hypothesis that has not previously been recovered cladistically and is also not supported by the existing taxonomic literature of dicynodonts. Ultimately, the repeated overestimates for clade divergence dates are driven by older taxa and sometimes entire clades being recovered as nested in younger clades, usually what is expected to be their sister group.

For origin times, the discrete and combined multirate models produce the most reasonable results given prior work. While the other analyses produce results that are somewhat older (Fig. 2a), they still have considerable overlap. The more striking concerns in the taxon ages are exemplified in the temporal disconnect between the two species of *Eodicynodon*. Although both “*Eodicynodon*” (now *Nyaphulia*) *oelofseni* and *Eodicynodon oosthuizeni* were found at the same locality, mere meters away from one another, only the combined-multirate analysis recovers these species as synchronous. This is most likely due to the ways in which we collect paleontological data, and the lack of autapomorphic characters in standard phylogenetic inference in paleontology. Because true autapomorphies, traits that are only present in terminal tips, do not influence the results of a parsimony analysis, they are generally described within the text, but very seldom are they coded into and included in the phylogenetic character matrix. This lack of coded autapomorphies means that the characters that differentiate tips are dominated by reversals or local autapomorphies. In the case of *Eodicynodon*, in our temporally distinct analyses we recover *E. oosthuizeni* as the sister taxon to *Lanthanostegus mohoii*. This relationship is important because it would require multiple reversals of synapomorphies in *E. oosthuizeni* to produce such a relationship. The additional local autapomorphies in *E. oosthuizeni* offer an explanation for the differing ages reconstructed in otherwise synchronous species. It is likely that the inclusion of additional autapomorphies would help provide better age constraints in Bayesian analyses (Matzke and Irmis, 2018). The most important part of our models are their failures, which make perfect sense given the nature of paleontological data and the present limits around using character evolution models in phylogenetic estimation. Perhaps the most important note for paleontologists is to include autapomorphies in character matrices; to not only incorporate autapomorphies into the same matrices as, and context in which we identify synapomorphies, but also to allow for future analyses and comparisons between parsimony and tip-dated phylogenetic inference. This is, additionally, an interesting consideration for methods developers to consider the role of lagerstattë effects in the FBD model.

### 4.4 An objective future for fossil phylogenetics

Molecular phylogenetics has long established incorporating multiple forms of information for joint estimation, sometimes labeled as ‘total evidence’ analysis. Over the years this has often incorporated both molecular, morphological, and sample age evidence (Heath et al., 2014). But it has more seldomly included continuous trait evidence (Parins-Fukuchi, 2018; Álvarez-Carretero et al., 2019; Zhang et al., 2024). In the field of comparative methods, there are established numerous continuous character evolution models, including Brownian motion and its variants, and the ways that they evolve along a phylogenetic tree. Joint inference using character evolution models is a critical step forward in Bayesian phylogenetics, as any number of data types can be included flexibly by expanding the model graph (Warnock and Wright, 2020). This allows continuous variables to be included and used to explore distinct regions of traitspace that may be otherwise uninvestigated. Uniting the simplest character evolution model (BM) along with the simplest morphological model (Mk, Lewis (2001)) establishes a critical starting point for advancing phylogenetic inference in the paleobiological sciences. Further questions, such as how character correlations and/or dependency affect topology, can be usefully asked in this framework.

Likelihood-based phylogenetics has historically attempted to incorporate continuous variables through a variety of models. These include the threshold models, which discretize continuous variables that have numerical values exceeding a threshold (Felsenstein, 1988), and Ornstein-Uhlenbeck models, which have trait optima (Hansen, 1997). These models have mostly been used for *post hoc* trait evolution inference. This represents some of the earliest work in incorporating character evolution models into the estimation of a phylogenetic tree (but see Parins-Fukuchi (2018); Zhang et al. (2024)) using a dataset for which both continuous and discrete traits are available, enabling direct signal comparison. In both of these studies, continuous traits were found to be a significant boon for phylogenetics, due to their relatively lower evolutionary rates and lack of homoplasy. While this study has highlighted some complications to the incorporation of continuous traits, we’ve also fundamentally found that these traits are contributing meaningful signal that we can model to improve our phylogenetic estimations.

The incorporation of continuous variables will be a net benefit to all life sciences, but will have a disproportionately large impact in paleontological communities, where information is limited due simply to the process of fossilization. These joint inference models provide flexibility in the kinds of data and the way those data are treated, which in turn provides the investigator more information in an otherwise limited framework. As model choice appears to have a very strong impact on tree inference and the ability to reach convergence, it is inevitable that additional character evolution models, such as the Ornstein-Uhlenbeck model which is meant to be an approximation of selection, will be necessary to obtain our best approximations of phylogeny. Additionally, among-character correlations may be an important next step in this work. Developmental processes and modular evolution (Clarke and Middleton, 2008) can cause suites of traits to be non-independent of one another, a fact which likely affects both discrete and continuous traits. This problem has received relatively little attention in Bayesian analyses, but may lead to an over-representation of some signals within the data.

The field of paleontology has a long history of co-opting neontological tools and ideas, usually a decade or two after original publication, for use with fossils. This has led to paradigm shifts within the field, including ecological abundance indices, CT-scanning, and in our opinion, Bayesian phylogenetics is likely to be a future paradigm shift in the field. Joint inference integrates neontological concepts and tools, such as clock models, with modern quantitative paleobiological tools, such as the fossilized birth-death process. The useful synthesis of paleontological and neontological ways of viewing the world has provided a unique framework for testing the impact of different character evolution models on phylogenetic estimation. Core issues remain, such as that the direct quantification of morphology tends to either obscure nuance when changes are textural, or they require high-resolution 3D scans accompanied by advanced computational methods that can produce and compare topological maps of morphology (Goswami et al., 2014; Fabre et al., 2020; Boyer, 2008). As we propose in section 4.1, there are numerous ways forward to deal with these issues. We hope this work will form a basis for interesting and productive work on the statistical problems associated with continuous trait incorporation in phylogenetic analyses.

## 5 Acknowledgments

AMW, BMW, and BK were supported on NSF DEB 2045842 and AMW was supported on NSF CIBR 2113425. AMW additionally received funds from Southeastern Louisiana University’s Dyson Fellowship. PJW was supported on NSF EAR 2129628. BK was additionally supported by the European Union (ERC, MacDrive, GA 101043187). Views and opinions expressed are however those of the authors only and do not necessarily reflect those of the European Union or the European Research Council Executive Agency. Neither the European Union nor the granting authority can be held responsible for them. Finally, we thank associate editor Dr. Joëlle Barido-Sottani, reviewer Dr. Caroline Abbott, and two anonymous reviewers for their incisive and helpful comments.

## 6 Data Availability

All the original data and scripts necessary to reproduce the analyses reported in this study can be accessed through the Dryad link: http://datadryad.org/share/ahyNsjmSGsFvdaQ0SvVhmUj2Jmp0hnhnlK8Wlp_ensQ

---

**Supplementary Figure S1:**
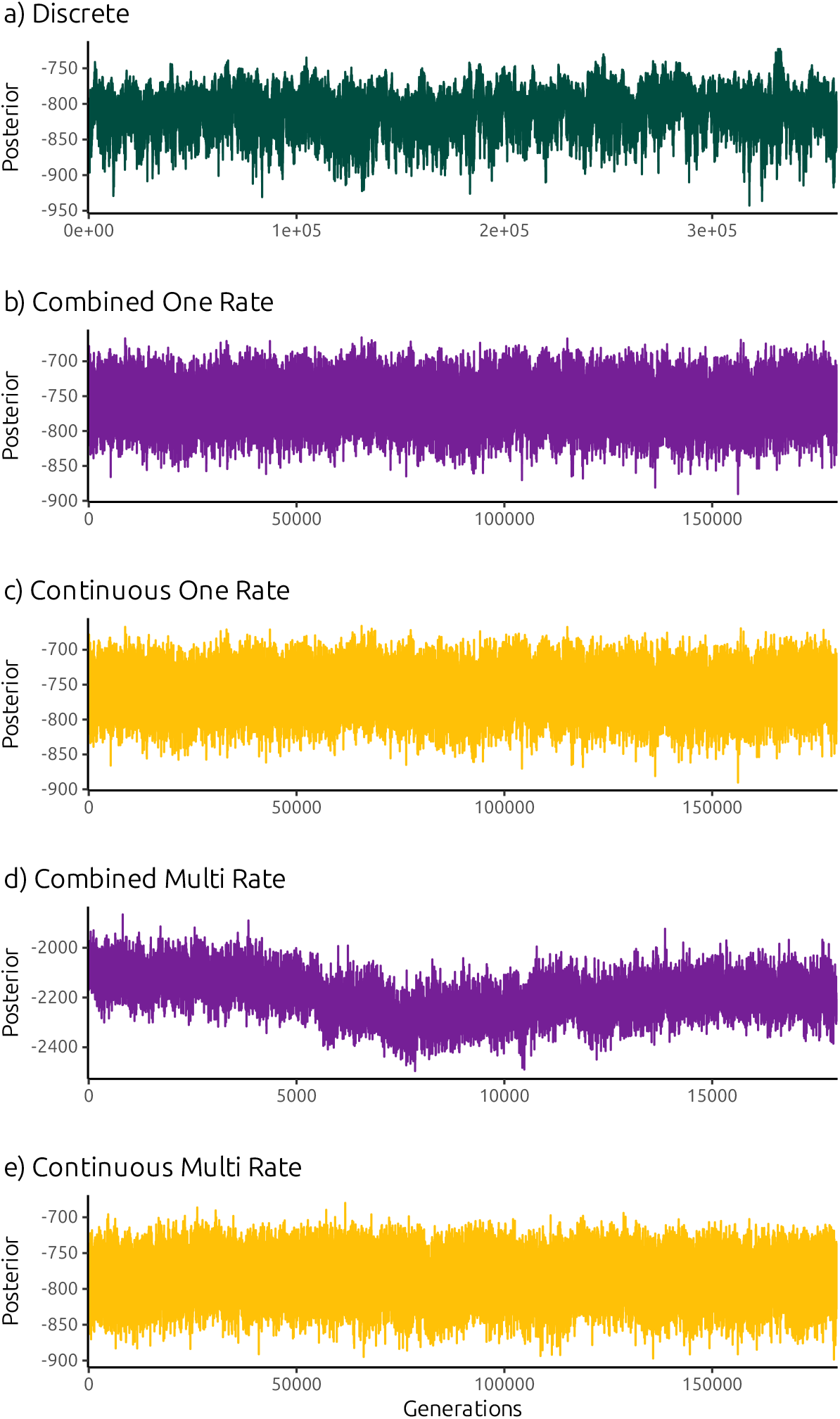
Trace plots of the posterior probability demonstrating that the analyses are converged.

**Supplementary Figure S2:**
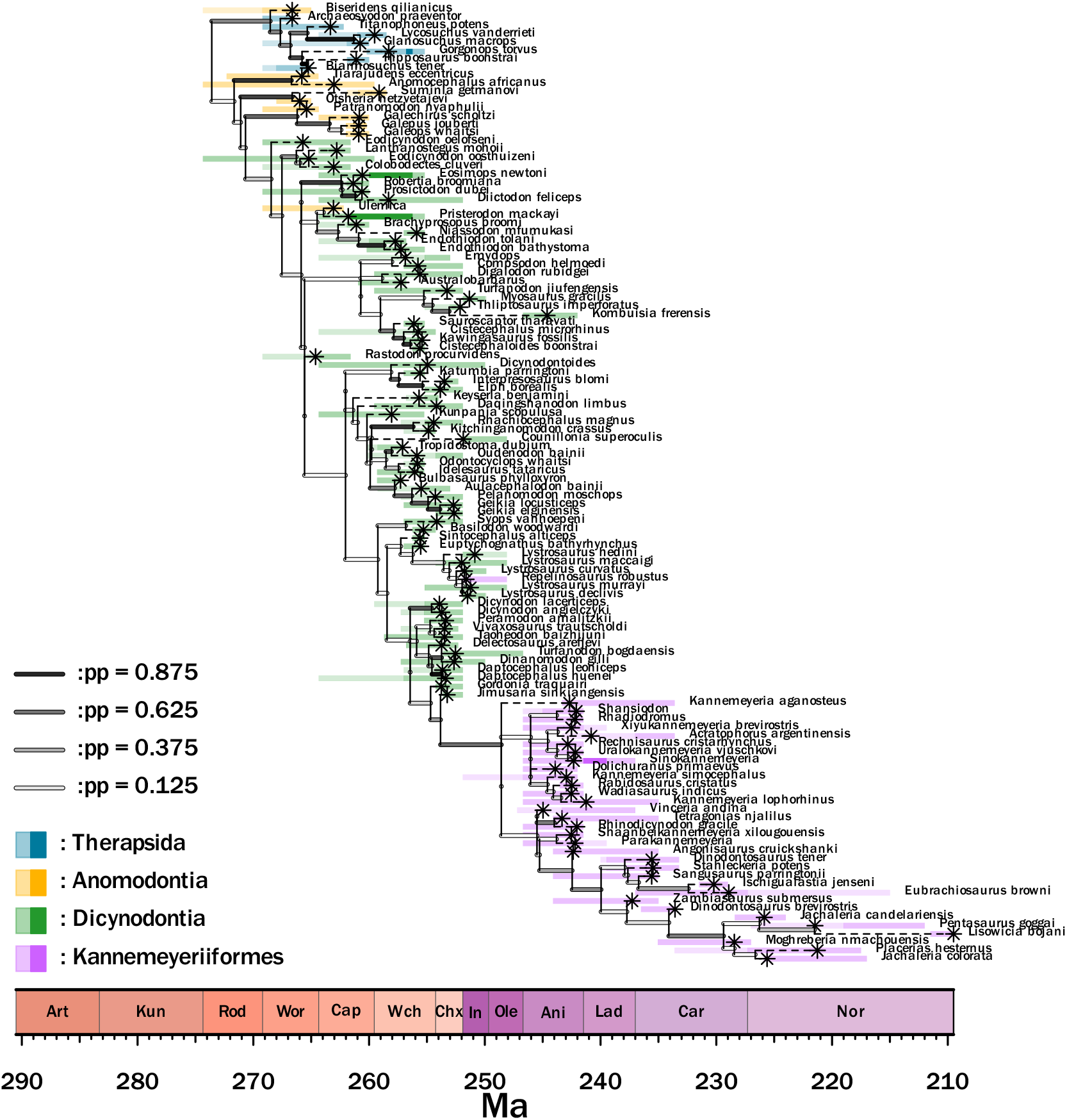
A dated phylogenetic tree estimated from discrete morphological character information.

**Supplementary Figure S3:**
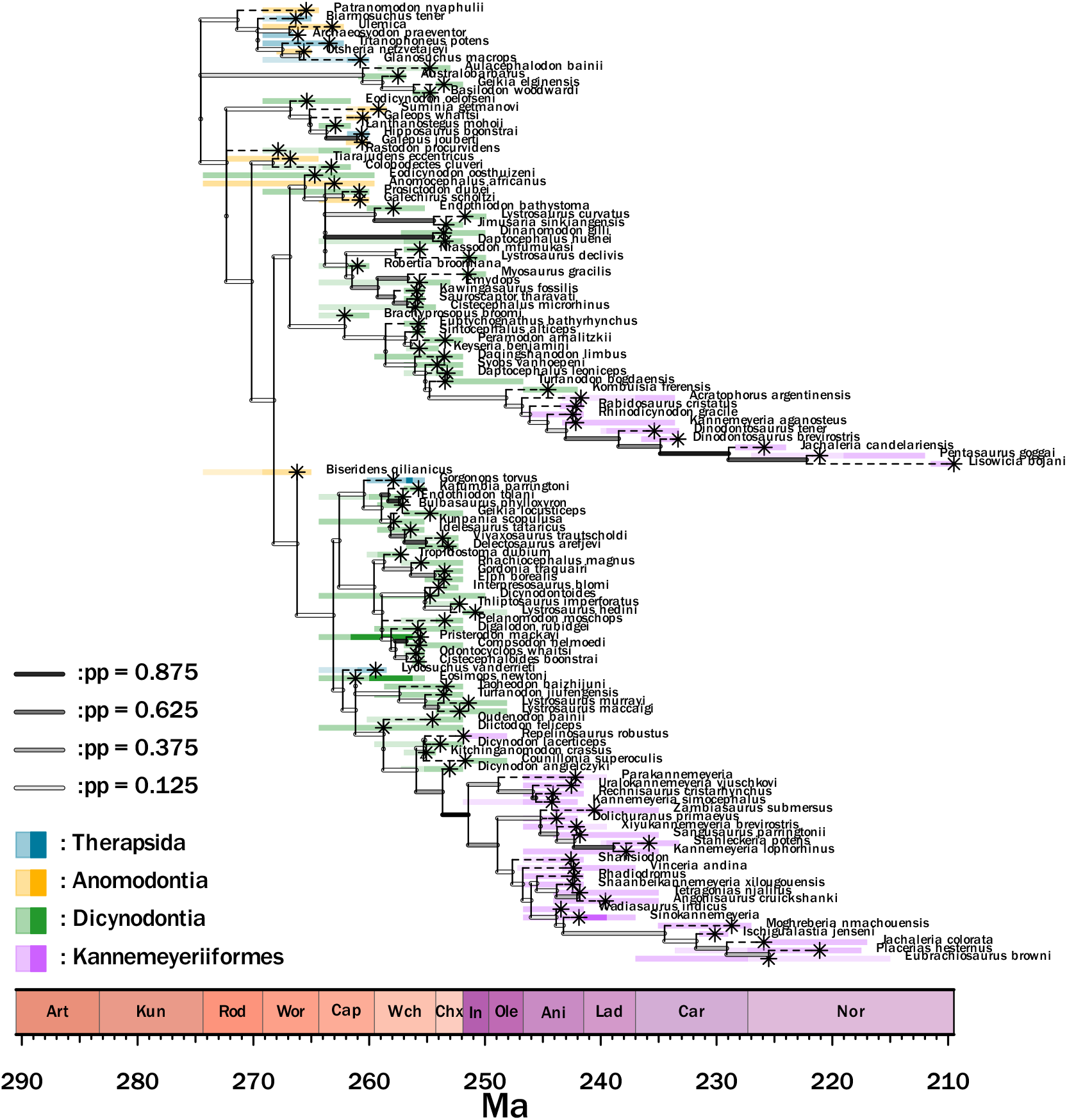
A dated phylogenetic tree estimated from continuous morphological character information. In this analysis, all continuous traits are assumed to share a single Brownian Motion model.

**Supplementary Figure S4:**
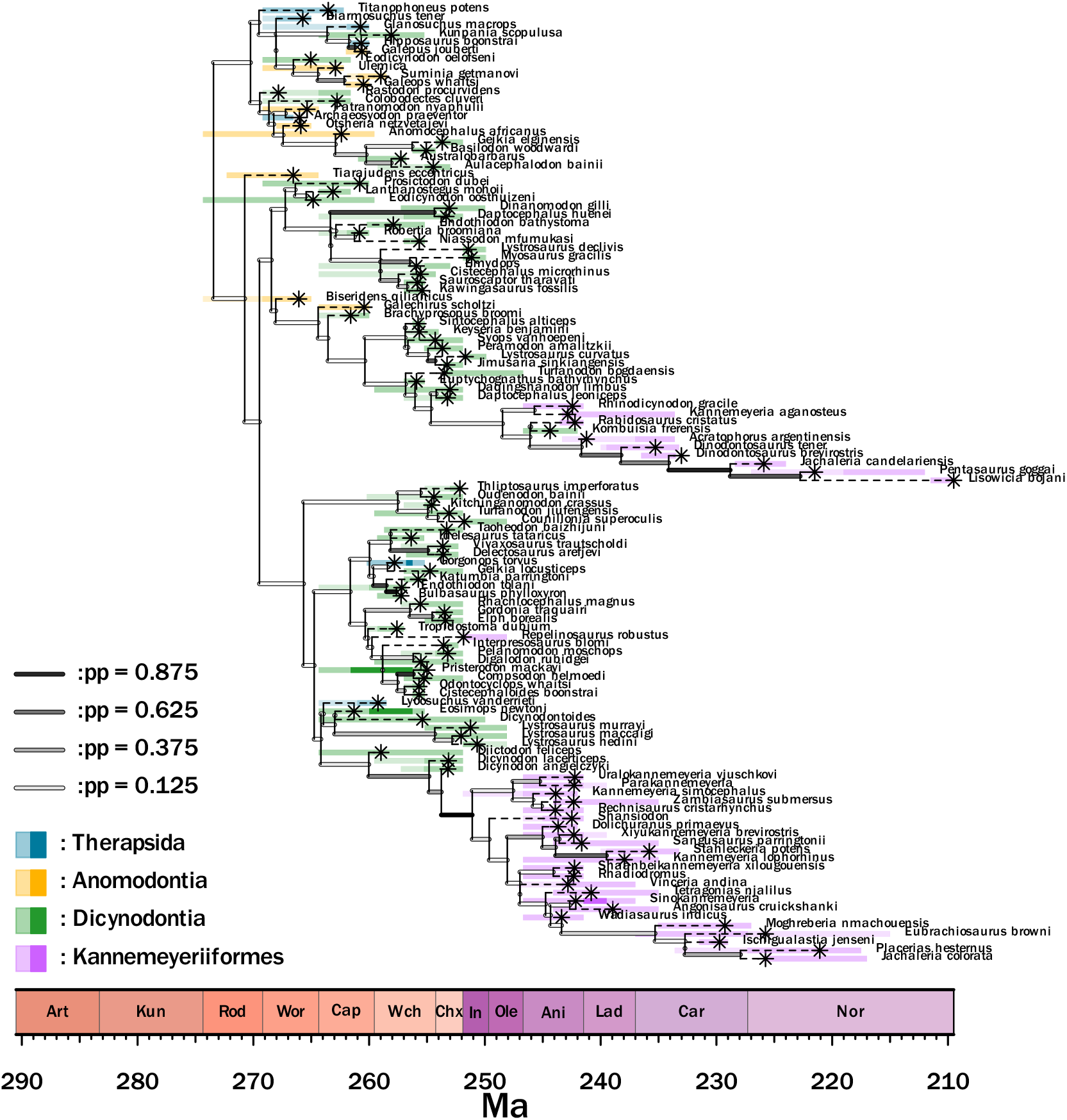
A dated phylogenetic tree estimated from continuous morphological character information. In this analysis, all continuous traits are assumed to have their own Brownian Motion model.

**Supplementary Figure S5:**
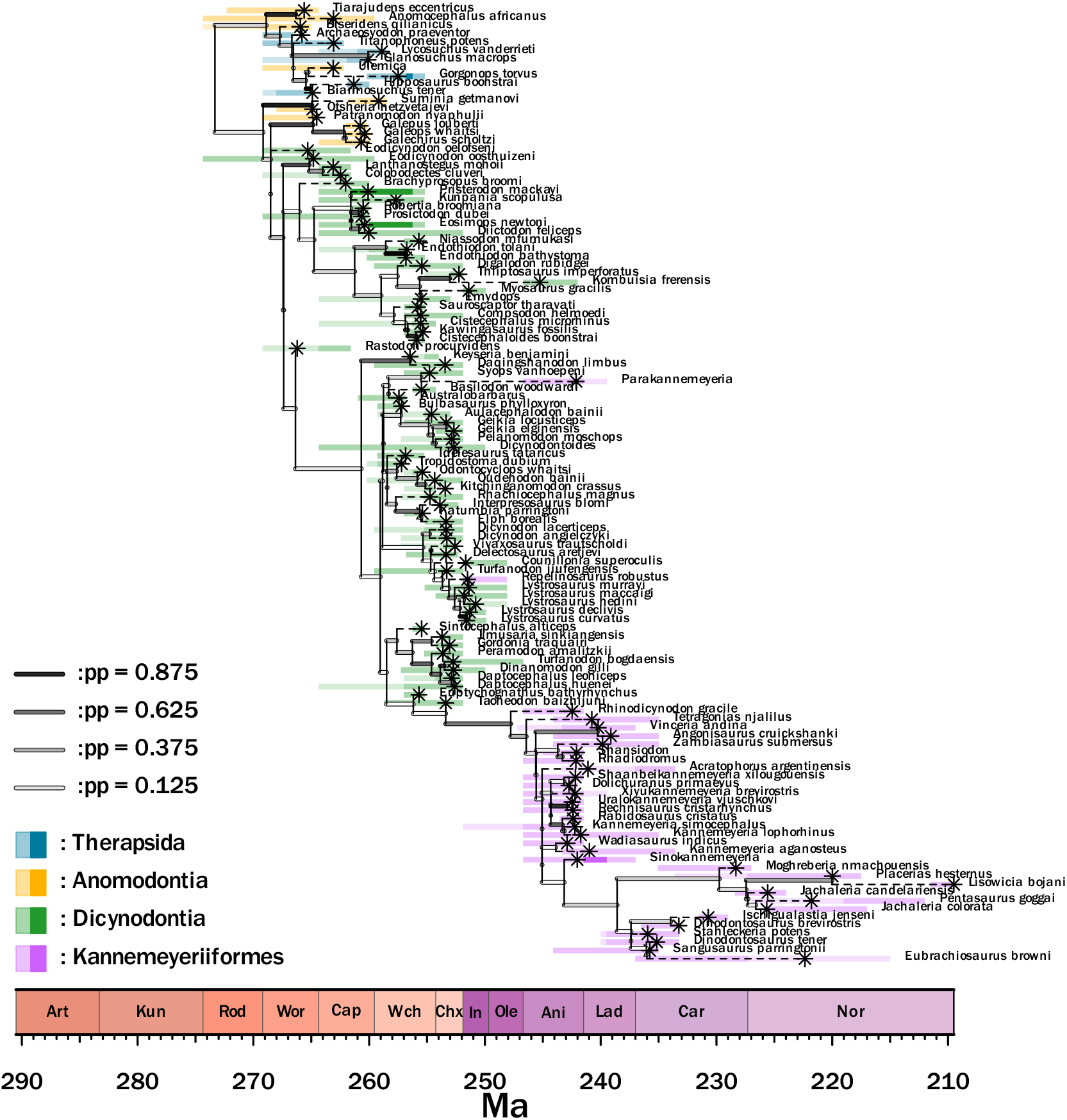
A dated phylogenetic tree estimated from continuous and discrete morphological character information. In this analysis, all continuous traits are assumed to share a single Brownian Motion model.

**Supplementary Figure S6:**
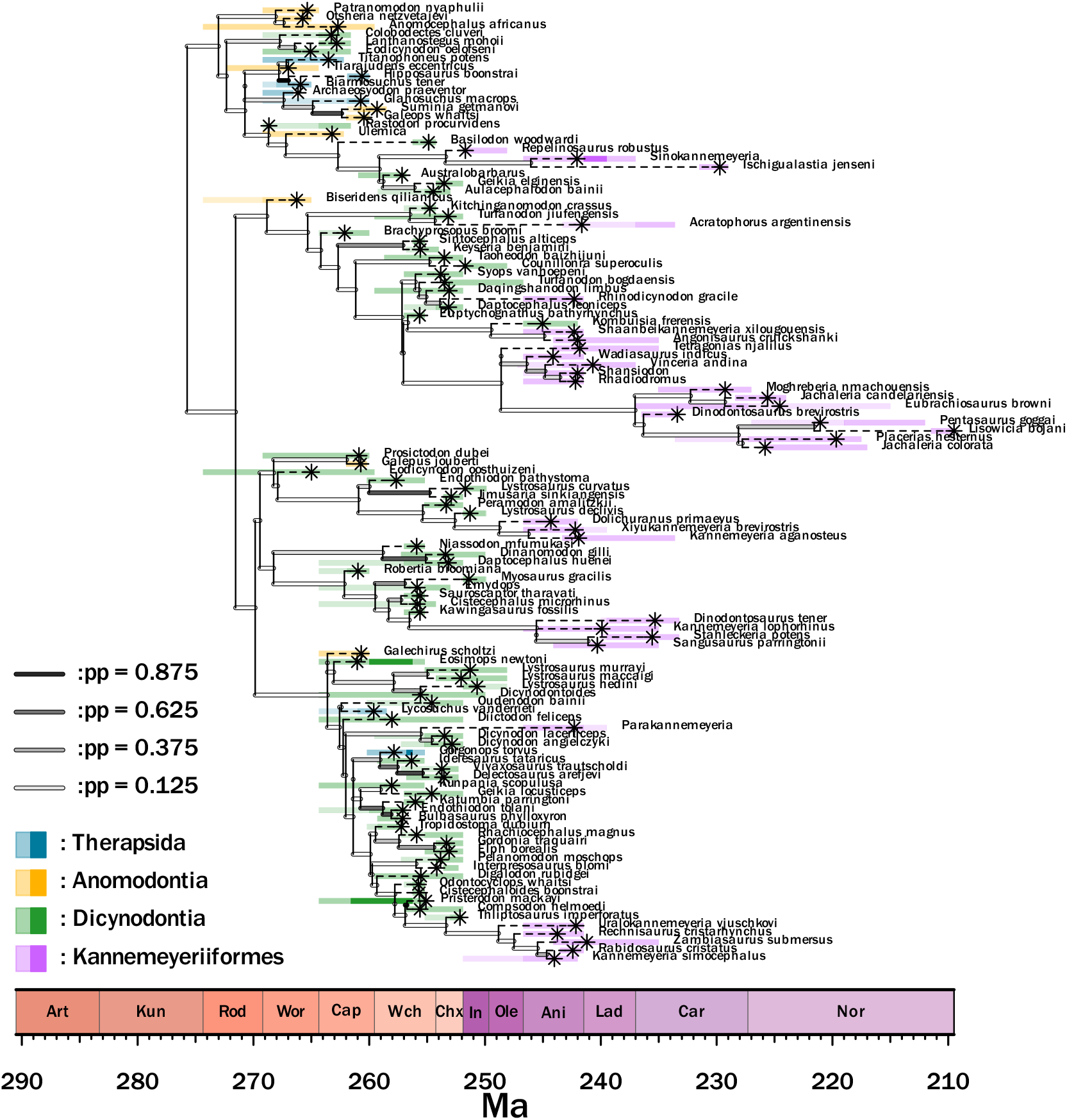
A dated phylogenetic tree estimated from continuous and discrete morphological character information. In this analysis, all continuous traits are assumed to have their own Brownian Motion model.

**Supplementary Figure S7:**
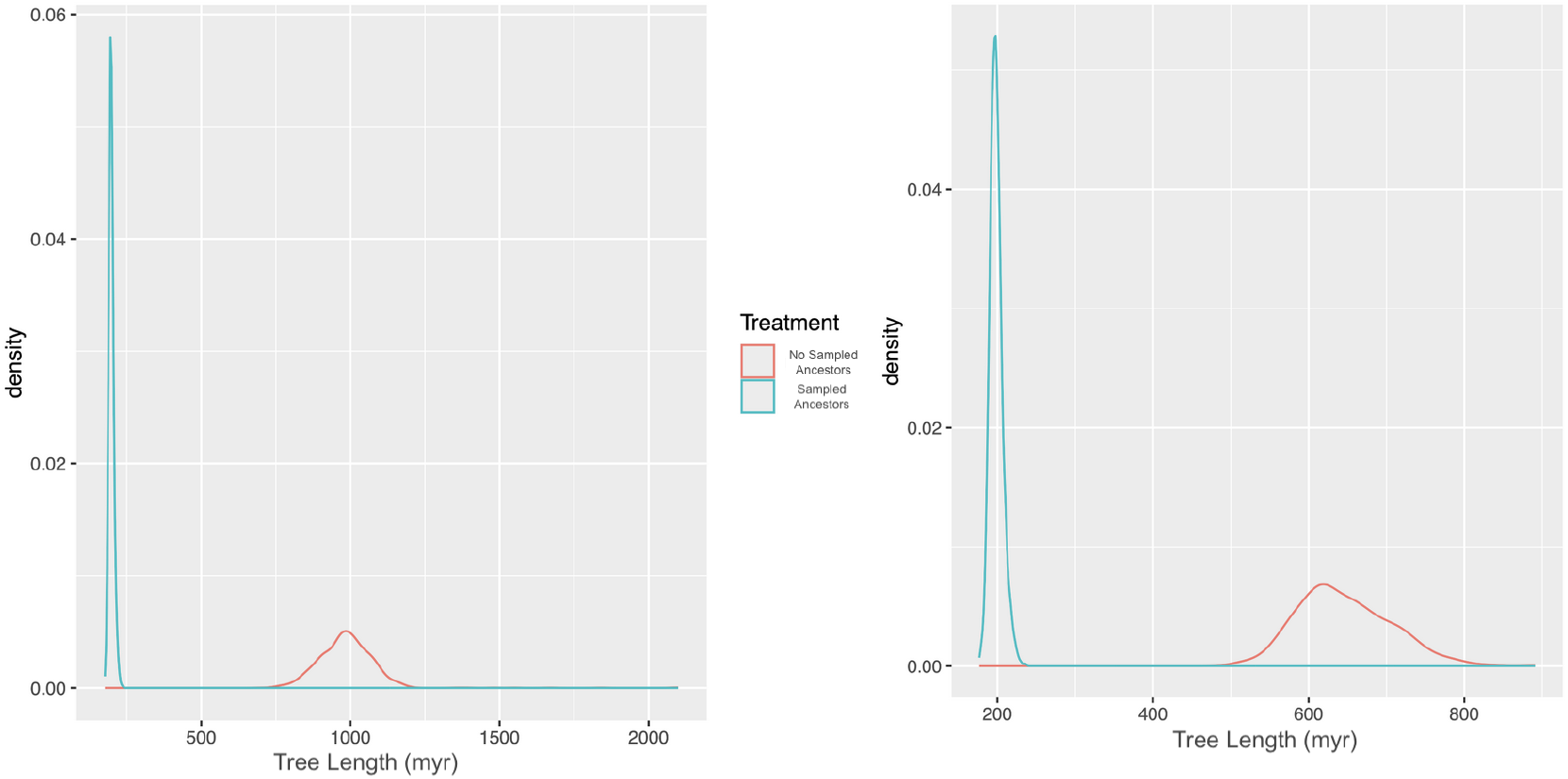
Two plots demonstrating the effect of including sampled ancestors on the analysis. At left, an analyses using both discrete and continuous data, with the continuous data assumed to share one Brownian Motion process. At right, the same data, but with a multi-rate Brownian motion process. In both, excluding sampled ancestors results in a significantly longer tree. However, including sampled ancestors does mean MCMC must be run longer due to the increased tree space size.

## Notes

http://datadryad.org/stash/share/ahyNsjmSGsFvdaQ0SvVhmUj2Jmp0hnhnlK8Wlp_ensQ

### Competing Interest Statement

The authors have declared no competing interest.

### Summary of Updates

New figures at the request of reviewers. Improved figure resolution.

